# Caffeine-tolerant mutations selected through an at-home yeast experimental evolution teaching lab

**DOI:** 10.1101/2023.01.17.524437

**Authors:** Naomi G. Moresi, Renee C. Geck, Ryan Skophammer, Dennis Godin, yEvo Students, M. Bryce Taylor, Maitreya J. Dunham

## Abstract

yEvo is a curriculum for high school students centered around evolution experiments in *S. cerevisiae*. To adapt the curriculum for remote instruction, we created a new protocol to evolve non-GMO yeast in the presence of caffeine. Evolved strains had increased caffeine tolerance and distinct colony morphologies. Many possessed copy number variations, transposon insertions, and mutations affecting genes with known relationships to caffeine and TOR signaling - which is inhibited by caffeine - and in other genes not previously connected with caffeine. This demonstrates that our accessible, at-home protocol is sufficient to permit novel insights into caffeine tolerance.

## DESCRIPTION

yEvo (“yeast Evolution”) was developed as a curriculum for high school students to perform laboratory evolution experiments with *Saccharomyces cerevisiae* yeast in their classrooms (Taylor et al. 2022a). We wanted to expand the curriculum to include other selective pressures and make the yEvo experiments safe and accessible to instructors and students in a variety of classrooms and for remote learning. The first iteration of this curriculum evolved azole-resistant yeast using a common laboratory strain (Taylor et al. 2022b), but as a genetically modified organism (GMO), this strain is not suitable for all instructional settings. Transition to remote learning during the COVID-19 pandemic also led to creation of resources for at-home experiments, including kits for undergraduate students to evolve resistance to household cleaners in store-bought baking yeast (Bennett et al. 2021). For our experiments with high school students, we used Fleischmann’s baking yeast, a non-GMO *S. cerevisiae* strain available at grocery stores, and non-toxic chemicals for the selective pressure so that these experiments could be conducted outside the lab.

We chose caffeine as our selective pressure since it is relatively safe and has broad applications: *S. cerevisiae* is used to ferment foods containing caffeine, and studying caffeine tolerance increases our understanding of TOR (target of rapamycin) signaling since Tor1 is inhibited by caffeine (Ruta and Farcasanu 2020). TOR signaling is a nutrient-sensitive pathway that plays a role in cell growth, metabolism, and aging, and has many conserved functions across eukaryotes (González and Hall 2017, Morozumi and Shiozaki 2021). Caffeine inhibits TOR signaling and growth through interactions with the rapamycin-binding domain, compromising overall cellular fitness (Reinke et al. 2006). While many genes that regulate the TOR signaling pathway are known, there may be more to discover. Furthermore, how other pathways compensate for TOR signaling inhibition is not completely understood (Ruta and Farcasanu 2020).

Our goal was to work with high school students to identify genes involved in TOR signaling. We did this by growing yeast in inhibitory concentrations of caffeine to select for better-growing mutants with increased caffeine tolerance. Since the Fleischmann’s yeast strain is one of the most sensitive to caffeine within a collection of 1,011 diverse *S. cerevisiae* isolates (top 25) (Peter et al. 2018, Schacherer, personal communication), we hypothesized that we could evolve caffeine tolerance in Fleischmann’s yeast in a relatively short period of time amenable to integration into a high school biology class.

During the shift to remote instruction in the COVID-19 pandemic, 28 students were given supplies to evolve Fleischmann’s yeast for eight weeks at home in increasing concentrations of caffeine. These at-home evolution kits included a starter culture of Fleischmann’s yeast, disposable test tubes containing a range of concentrations of caffeine in an acidified media (pH 4.6) to reduce risk of bacterial growth, and sterile swabs (see methods for more details). Students’ evolved yeast were then sent to the University of Washington, where clones were isolated from each evolved population for whole-genome sequencing and phenotyping. Some of the students reported bacterial contamination during their evolution experiments, but all isolated evolved clones were confirmed to be *S. cerevisiae* during sequencing.

Clones isolated from evolved populations had increased caffeine tolerance, as they grew at concentrations of caffeine above 6.25mM, which inhibited growth of the ancestral strain (Fig. 1A). When grown on YPD agar plates without caffeine, evolved clones also exhibited different colony morphologies. Many formed petite colonies, which have been previously linked to increased caffeine tolerance (Bard et al. 1980). Most evolved clones lost their ancestral wrinkled colony morphology (Fig. 1B), which correlated with their increased caffeine tolerance (Fig. 1A). Since many genes that contribute to wrinkled colony morphology and filamentous growth have also been shown to be required for flocculation (Lo and Dranginis 1998, Reis 2014), we also tested the effect of caffeine on flocculation. We observed that the ancestral strain and 5/6 evolved clones that formed wrinkled colonies also flocculated, while clones with smooth morphology did not (Extended Data Table 1). These morphological changes could have industrial consequences since flocculation can affect industrial fermentation processes, and health applications since filamentous growth in *S. cerevisiae* can be used to model virulence of the pathogenic yeasts, as reviewed in Brückner et al. 2012. Further experiments will be required to determine if these morphology changes are directly causal for improved caffeine resistance and/or improve successful transfer in our serial culture procedure.

**Figure 1:**
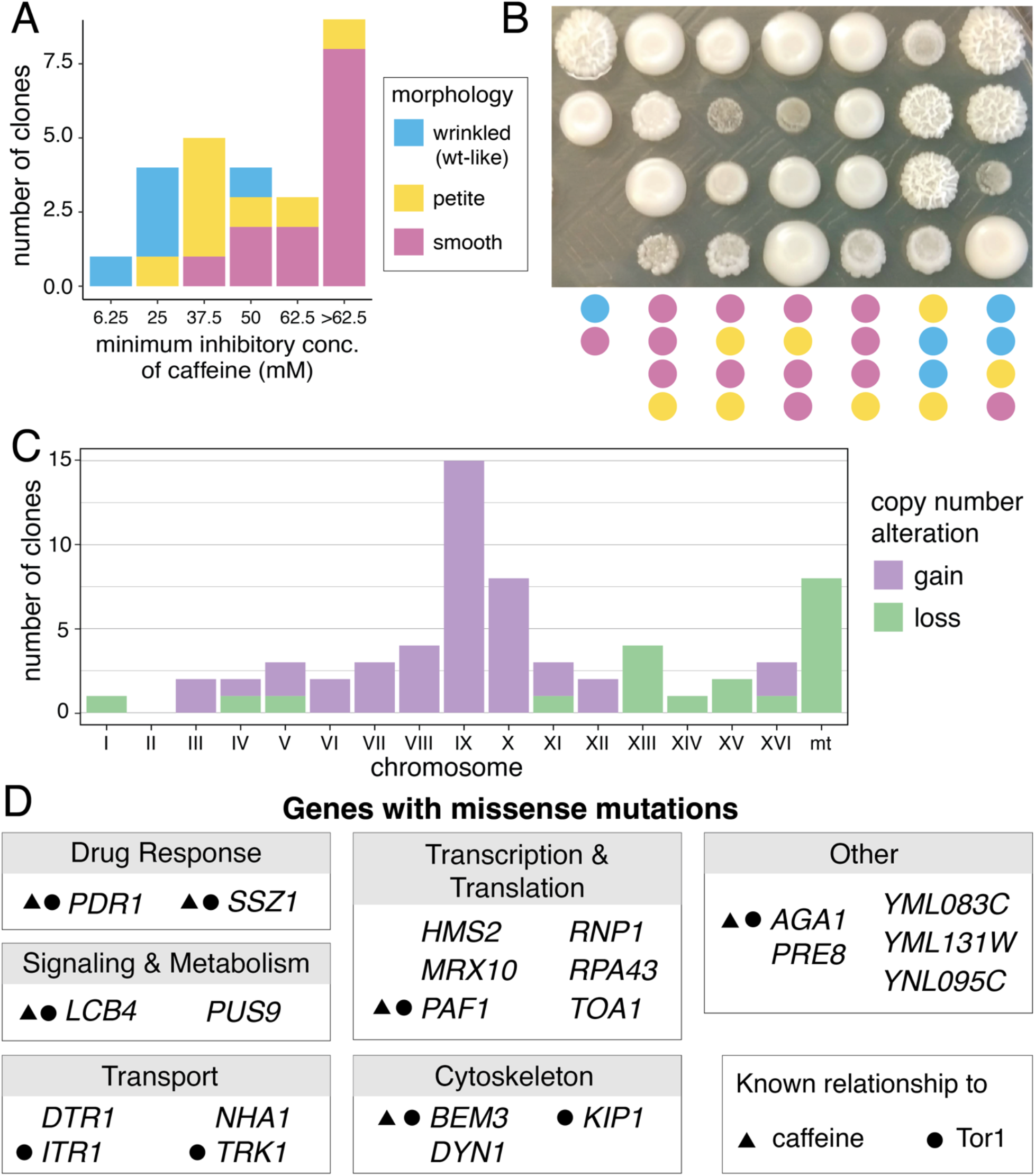
Characteristics of caffeine-tolerant Fleischmann’s yeast. A. Minimum inhibitory concentration of caffeine for clones isolated from caffeine-evolved populations. B. Representative colony morphologies of evolved clones. Morphology classification indicated by color scale below image and legend in A. C. Number of instances of chromosomal gains and losses in evolved clones, compared to 2N euploid ancestor. All gains indicate 3N, and losses represent 1N except mitochondrial DNA (mt) loss which is a complete loss of mitochondrial chromosome. D. All genes with nonsynonymous mutations found in caffeine-evolved clones. Genes grouped by function from SGD. Symbols denote previous publications associating the gene with caffeine or Tor1. All publications are cited in the main text.

We performed whole-genome sequencing on 26 clones from 22 caffeine-evolved populations. The majority (21/26) of caffeine-tolerant clones had at least one whole chromosomal gain or loss, compared to their ancestral 2N euploid state (Fig. 1C and Extended Data Table 1) (Peter et al. 2018). Eight clones lost their mitochondrial DNA, which leads to their petite phenotype and confers resistance to many drugs, including caffeine, through increased activity of the pleiotropic drug response network (Bard et al. 1980, Hallstrom and Moye-Rowley 2000). 8/26 clones gained a copy of chromosome X, which contains the *TOR1* gene and thus may confer some resistance to Tor1 inhibition. The most frequently gained chromosome (15/26) was chrIX, which has been gained in other drug-adapted yeast populations (Chen et al. 2015) and has loci associated with rapamycin response (Lin et al. 2018). Only one evolved clone had a transposon insertion (Extended Data Table 1): a heterozygous Ty2 insertion upstream of *RNR4* and *TIM13*, which have both been implicated in response to caffeine and rapamycin (Xie et al. 2005, Kapitzky et al. 2010, Nicastro et al. 2021).

We identified non-synonymous mutations in 13/26 evolved Fleischmann’s yeast clones (Extended Data Table 2). All mutations were heterozygous, with the exception of the mutation in *RPA43*, which was found in two clones from the same population: homozygous in one clone and present in a single copy in the other due to loss of chromosome XV. Some other mutations were on chromosomes that were gained, but it was not possible to definitely determine if the mutant or wildtype copy was gained based on variation in read coverage. We did not find an association between any specific mutations or copy number changes and the morphology phenotypes. The changes in morphology may thus be due to mutations we were unable to detect by short read sequencing, by a combinatorial effect of multiple mutations and copy number variations (e.g. Tan et al. 2013), or by nongenetic changes, such as in prions, which have previously been shown to cause colony morphology changes (True and Lindquist 2000, Holmes et al. 2013).

Genes containing mutations in caffeine-evolved clones are involved in processes such as translation, transcription, signaling, and metabolism, all of which can be affected by caffeine and TOR (Fig. 1D and Extended Data Table 2). In both clones isolated from one population, we found a mutation in *PDR1*, which has been shown to contribute to caffeine tolerance in a previous experimental evolution study (Sürmeli et al. 2019). We also identified missense mutations in genes previously found through deletion set screens to decrease caffeine and rapamycin tolerance (*BEM3, LCB4, PAF1, PDR1, SSZ1, TRK1*) (Betz et al. 2002, Parsons et al. 2004, Dudley et al. 2005, Xie et al. 2005, Brown et al. 2006, Banuelos et al. 2010, Kapitzky et al. 2010, Nicastro et al. 2021). Other genes with mutations have been connected to caffeine and TOR signaling through studies in yeast (*ITR1*) (Hatakeyama and De Virgilio 2019) or of homologs in other organisms (*KIP1*) (Jang et al. 2020). Additionally, there were missense mutations in 13 genes not previously connected to caffeine or TOR, which merit additional research to identify the mechanisms underlying their relationships. We anticipate that many of these will be neutral as most evolution experiments find a large proportion of neutral mutations (Payen et al. 2016, Buskirk et al. 2017), but further investigation could reveal that some are connected to caffeine tolerance and TOR signaling. Every clone had at least one nonsynonymous mutation or copy number variant, so there are candidates for genetic contribution to caffeine tolerance in every clone. Only two genes were found mutated in more than one independent population (*TRK1* and *NHA1*, each in two populations). These two genes may be selected for by growth in the acidified media since they encode ion channels and contribute to acid tolerance (Kawahata et al. 2006, Mira et al. 2009). Additional replicate populations performed by future classes could help identify repeated targets of selection that would help highlight which candidates are most likely to be causative.

Taken together, our results showcase a protocol that makes experimental evolution more accessible for the classroom, remote learning, and homeschooling. The Fleischmann’s yeast strain was safe to use outside of a classroom or laboratory, and enabled students to visually observe phenotypic changes as over time they could see more yeast growing in tubes with concentrations of caffeine that inhibited growth of the ancestor. In the future, we would like to develop a protocol for students to observe the changes in colony morphology. We would additionally like to include more opportunities to directly measure increased caffeine tolerance at home, such as by conducting the minimum inhibitory concentration assays we performed in the laboratory, or by disk diffusion assay as used in other at-home evolution strategies (Bennett et al. 2021). Through this experiment, high school students contributed to ongoing research and their work may lead to the discovery of novel roles for genes in caffeine tolerance and TOR signaling.

## METHODS

### Experimental evolution

Students were given disposable sterile plastic test tubes (Corning 352059) containing a suspension of Fleischmann’s yeast in media containing 10mM caffeine. Yeast were cultured in YPD acidified to pH 4.6 (29.9g sodium citrate, 12.4mL 13M hydrochloric acid, 10g yeast extract, 20g peptone, and 20g dextrose per liter) to reduce risk of contamination. Students used a sterile cotton swab to transfer part of their culture into a new tube containing YPD with caffeine one to two times per week for eight weeks. Students transferred to increased concentrations of caffeine (20mM and then 40mM) at their discretion; not all experiments reached the same final concentration. If a culture was contaminated with bacteria (observed to be cloudy), students performed a transfer from the preceding culture. At the end of the experiments, students returned tubes containing their evolved populations to their instructor, who prepared a glycerol stock from each final population. Upon receipt of evolved populations at the University of Washington, single colonies were isolated on YPD-agar plates and saved for phenotyping and sequencing. In sample names, letter codes denote different populations, and numbers differentiate clones from the same population.

### Safety

Students were instructed to dispose of swabs used to transfer yeast in household trash, since Fleischmann’s yeast is a non-hazardous household substance. All tubes containing yeast cultures were sealed and returned to the instructor in case of unidentified contamination.

### Minimum inhibitory concentration (MIC) assay

Clonal cultures were grown to saturation in acidified YPD, and 2μL was transferred to 200μL of acidified YPD containing 0-62.5mM caffeine in a 96-well plate. Presence or absence of growth was recorded after 48h at 30°C. Reported MIC (Extended Data Table 1) is the median of three technical replicates per clone.

### Colony morphology assessment

Clonal cultures were grown to stationary phase in 200μL YPD in 96-well plates and pinned onto YPD-agar plates. Colonies were photographed after 48h growth at 30°C. All petite colonies were confirmed to be ρ^0^ by inability to grow on YPG-agar plates and absence of mitochondrial DNA in whole-genome sequencing (Extended Data Table 1).

### Whole-genome sequencing

Sample preparation, sequencing, determination of copy number variation, and identification of SNPs and indels was performed as previously described (Taylor et al. 2022b). Briefly, DNA was purified using a phenol-chloroform methodology. This DNA was tagmented using a modified Illumina Tagmentation protocol (Baym et al. 2015). Strains were sequenced to an average coverage depth of 59 reads (range 30-105). Variants present in the ancestral strain were removed from consideration. Transposable element insertions were identified using McClintock v.2.0.0 (Nelson et al. 2017), filtered for non-ancestral insertions, and inspected if they were identified by at least three component methods per clone. All mutations and transposable element insertions were manually inspected in the Integrative Genomics Viewer (Robinson et al. 2011) to ensure they were not present in the ancestor and to determine zygosity. Mutations, transposon insertions, and copy number alterations for each clone are listed in Extended Data Tables 1 and 2. Annotation of gene function was based on Saccharomyces Genome Database gene description (yeastgenome.org). Sequencing reads are deposited in the NCBI Sequence Read Archive (SRA) under BioProject PRJNA910188.

## REAGENTS

| Strain | Genotype | Source/Reference |
| --- | --- | --- |
| BTC isolate 79 | Fleischmann's baker's yeast | Peter et al. 2018 |

## EXTENDED DATA

**Extended Data Table 1: Phenotypes, copy number variation, and transposon insertions in evolved clones.** Morphology, caffeine MIC, petite status, chromosomal gains and losses, and transposable element insertions in caffeine-evolved clones.

**Extended Data Table 2: Point mutations in evolved clones.** List of SNP and indel mutations in caffeine-evolved clones.

## ACKNOWLEDGMENTS

We thank Noah Fredstrom for assistance with sequencing, and Leah Anderson for mentoring and feedback. We thank Joseph Schacherer for personal communication of data related to caffeine tolerance of strains in Peter et al. 2018. We thank Clara J. Amorosi, Anton A. Adedipe, Leo A. Adedipe, Sayeh Gorjifard, Christopher R.L. Large, Rebecca Martin, Anja R. Ollodart, Sophia Showman, Soyeon Showman, Christine Queitsch, Margaux Walson, Taylor Wang, and Chiann-Ling Cindy Yeh for piloting the home evolution kits during the spring of 2020. We thank the following students for conducting selection experiments: Priya Agarwal, Tiffany Chang, Jessica Chen, Sophie Cheung, Emily Crowell, Isabella Delgado-Rojas, Riya Duddalwar, Sherry Fan, Hanna Goldberg, Amelia Hemmings, Millie King, Maya Lin-Stevens, Mirelle Linquist, Isabella Man, Maya Melnik, Xochitl Munoz, Jacqueline Pearce, Sara Sanchez-Lowin, Isabella Santoro, Samanta Scroggie, Katelyn Sim, Caledonia Steltzner, Jessica Tanouye, Genevieve Timoner, Sarah Tyebkhan, Katrina Weng, Sarah Yuhan, Charlotte Zhang.

